# Cognitive tests that identify high risk of conversion to dementia in Parkinson’s disease

**DOI:** 10.1101/2020.05.31.126870

**Authors:** D.J. Myall, K-L. Horne, M.R. MacAskill, L. Livingston, T.L. Pitcher, T.R. Melzer, G.J. Geurtsen, T.J Anderson, J.C. Dalrymple-Alford

## Abstract

**Background:** People with Parkinson’s disease who meet criteria for mild cognitive impairment are at increased risk of dementia. It is not known which tests are more effective than others for identifying the risk of dementia.

**Methods:** At baseline, we assessed performance on 21 neuropsychological test measures spanning five cognitive domains in a prospective longitudinal study of 196 non-demented people with Parkinson’s. Elastic net logistic regression was used to identify a pair of tests from each cognitive domain that best predicted conversion to dementia over a four year period. The optimal tests most predictive of dementia were also determined when mild cognitive impairment was derived from a logistic-regression classifier that used all 21 measures simultaneously.

**Results:** With two tests per domain, the resulting mild cognitive impairment group (N=87/196) captured 44 of 51 individuals who converted to PDD; the out-of-sample relative risk of PDD was 8.0 (95% CI [4.3, 24]), similar to that achieved with the full battery (N=102/196, capturing 45/51, relative risk = 6.9). When selecting tests regardless of domain, there was strong evidence for three tests: Trail Making part B (Executive), Map Search (Attention), and CVLT-II word list acquisition (Episodic Memory). The logistic-regression classifier achieved an out-of-sample AUC of 0.90 [0.84, 0.96] and a relative risk of 12 [6, 39].

**Conclusions:** An abbreviated selection of neuropsychological tests can identify non-demented patients who have a high relative risk of progression to PDD.

## Introduction

People with Parkinson’s disease (PD) often progress to dementia (PDD). Although some patients have cognitive impairment at the time of PD diagnosis, many only reach dementia after an extended period, if at all ^1–6^. Knowing the risk of PDD would clarify a patient’s prognosis, facilitate care and management, and enable suitably-powered clinical intervention trials ^7–10^. Recent work has focused on patients who meet the Movement Disorder Society Task Force (MDS-TF) criteria for PD with mild cognitive impairment (PD-MCI) ^11^. PD-MCI patients have an increased risk of progressing to PDD ^12–20^. Large multinational retrospective studies have confirmed that this risk is elevated beyond the influence of age, sex, education, motor severity and depression. This was found using both MDS-TF level I PD-MCI criteria, ^21^ with one test in each of five cognitive domains, and with level II criteria, which requires two tests in each domain ^22^. The risk of PDD in PD-MCI patients in these multinational studies was increased 2 to 3-fold for cognitive impairments that exceeded 1 SD below normative data and increased to 11 to 14-fold for cognitive impairments beyond 2 SD.

Significant cognitive decline and loss of independent function can develop relatively quickly in PD patients ^23,24^. Cognitive changes in those who progress to PDD suggest inflection points of accelerated decline two to five years prior to dementia diagnosis ^25,26^. Similarly, increased rates of conversion from PD-MCI to PDD have often been reported within three to five years after baseline assessment ^15,17–19,27^. Nonetheless, a meta-analysis of studies with follow-up of one to seven years reported that a considerable proportion of PD-MCI patients reverted to relatively normal cognitive function (PD-N) at both level II criteria (15%, 95% CI [11,21]%) and level I criteria (35%, [22,51]%); the latter rate was comparable to conversions to PDD (31%, [25,38]%) ^24^. Together, such evidence suggests that it is important to identify PD-MCI patients by focusing on baseline neuropsychological tests that are the most relevant for conversion to PDD within three to four years after cognitive assessment.

This goal requires direct comparison of a wide range of neuropsychological tests. Some studies suggest that frontal-executive impairments in planning and working memory are irrelevant for the risk of future dementia in PD ^28,29^. There is, however, mixed evidence both from before ^27,29–34^ and after the MDS-TF PD-MCI criteria ^12,15,16,18,35–39^ whether cognitive impairments associated with frontal brain circuitry dysfunction or posterior circuitry dysfunction provide optimal predictors of PDD risk. Many of these longitudinal studies had relatively small follow-up sample sizes or a relatively limited range of neuropsychological tests. Only two studies assessed the independent merits of different tests in PD-MCI patients ^16,39^. None have examined out-of-sample predictions for individual test measures.

We examined the value of 21 measures from sixteen neuropsychological tests as predictors of conversion to PDD in a large prospective longitudinal sample that was followed for four years after their baseline assessment. The tests spanned the five cognitive domains suggested by PD-MCI criteria ^11^. Our aim was to determine a reduced set of neuropsychological tests that would maximise the identification of a person’s imminent risk of PDD.

## Methods

### Participants

A convenience sample of patients meeting idiopathic UK Brain Bank PD diagnostic criteria ^40^ was recruited from an outpatient specialist movement disorders clinic and invited to participate in the ongoing New Zealand Brain Research Institute longitudinal study. Other movement disorders, a history of major developmental or adult neurological disorders or recent (6 months) major psychiatric disorders and evidence of PDD at baseline were exclusion factors. We followed 196 such individuals with PD who received cognitive assessments (median of every 2 years) for a period of between 3.5 and 4.5 years after their baseline assessment (Figure 1). The study was approved by the ethics committee of the New Zealand Ministry of Health and all participants provided informed consent.

**Figure 1:**
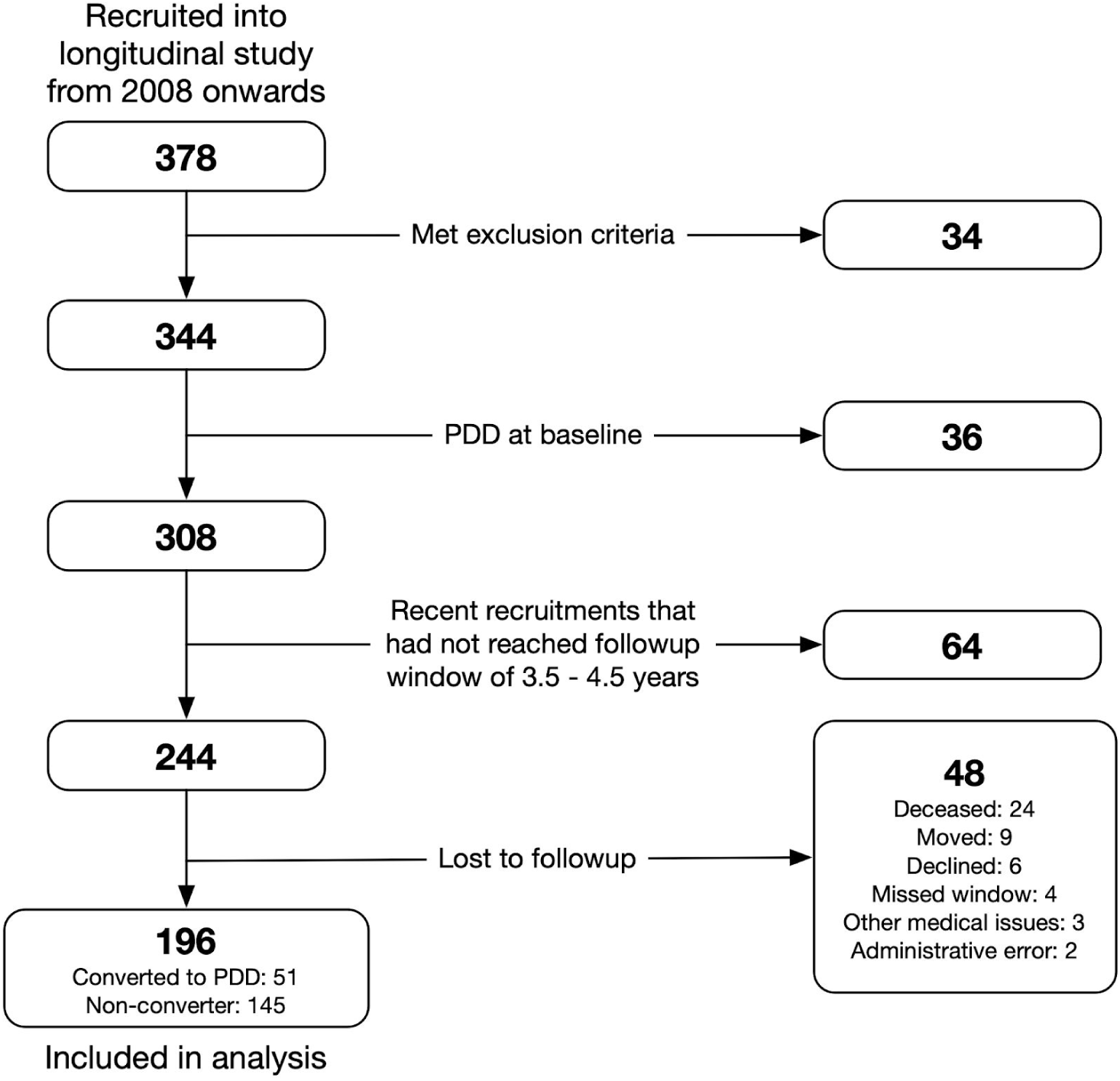
Flow diagram of participation

### PD-MCI and PDD diagnoses

Following the level II PD-MCI recommendations ^11^, we used neuropsychological tests that spanned five cognitive domains. These were administered and scored by trained research personnel over two sessions (Figure 2): (i) *Executive function* using Stroop interference, letter fluency, category fluency and category switching (all from the Delis-Kaplan Executive Function System (D-KEFS) ^41^), Trail Making part B, and action fluency; (ii) *Attention, working memory and processing speed*, using Map Search (first minute only; Test of Everyday Attention), digits forwards/backwards, digit ordering, Stroop color reading, Stroop word reading, Trail Making part A, and WAIS-III picture completion; (iii) *Episodic memory*, using the California Verbal Learning Test-II Short Form (CVLT-II SF) (total immediate recall across 4 trials; recall after 10-12 minutes), and uncued recall for the Rey Complex Figure Test (RCFT; at 3 minutes Immediate recall); (iv) *Visuoperception*, using judgment of line orientation (JLO), fragmented letters (Visual Object and Space Perception battery), and copy of the RCFT; and (v) *Language*, using the Mattis Dementia Rating Scale-2 (DRS-2) similarities components, and an aggregate score for the language components of the Alzheimer’s Dementia Assessment Cognitive Scale (ADAS-Cog, comprised of object and finger naming, commands, comprehension, spoken language and word-finding difficulties). Assessments also included neuropsychiatric measures (Neuropsychiatric Inventory; Geriatric Depression Scale; Hospital Anxiety and Depression Scale).

**Figure 2:**
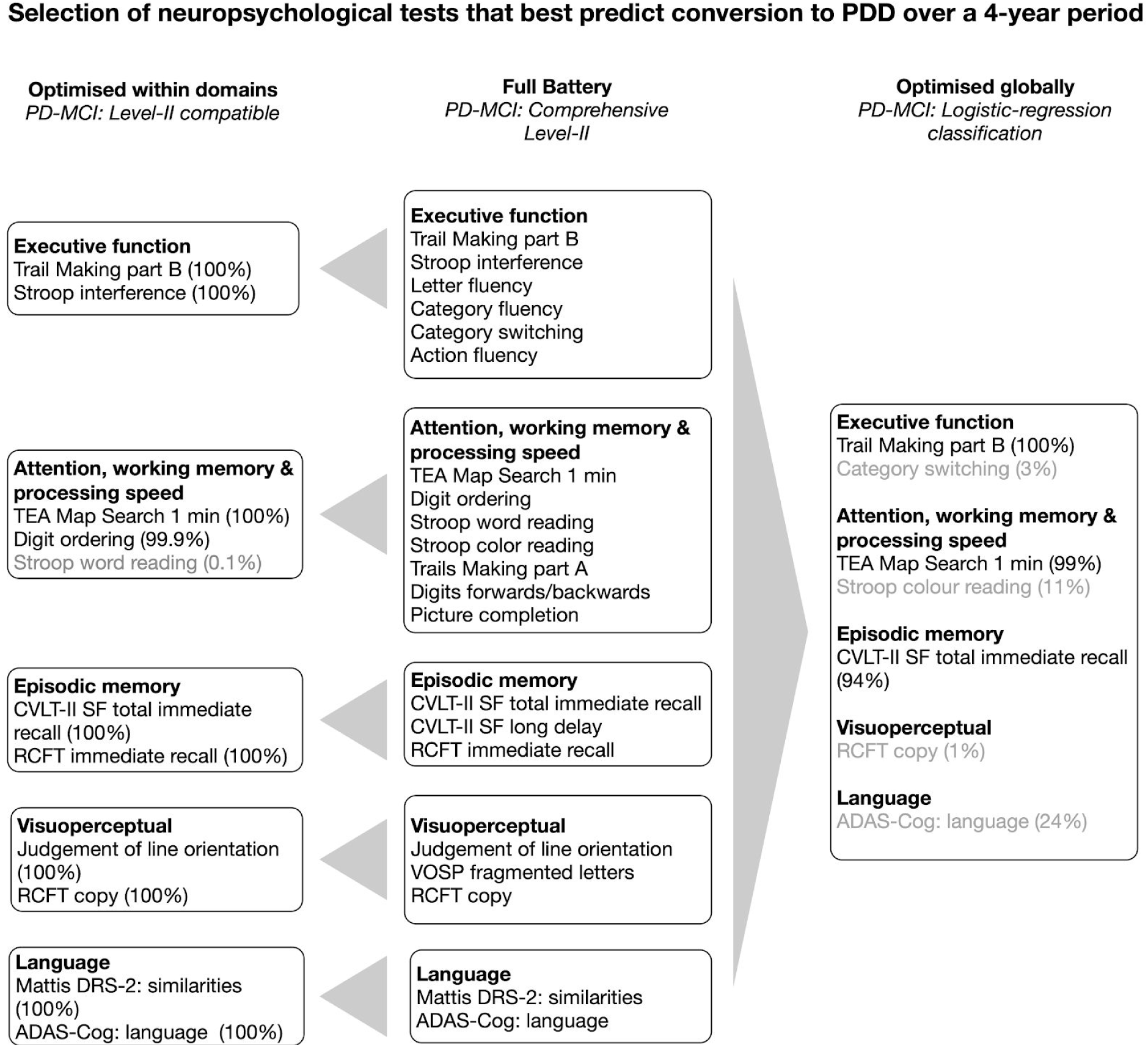
*Left*: tests identified when the analysis was constrained to select two tests per domain (percent = selection frequency over imputation and LOO loops). *Right*: tests identified when all 21 measures were entered simultaneously in a regularised logistic-regression classifier. *Centre*: the full list of neuropsychological tests.

A PDD diagnosis required both the presence of a substantial impairment (−2 SD or worse below normative data) in one or more tests in at least two of the five cognitive domains and evidence of significant decline from previous levels in everyday functional activities not attributed to motor impairments ^42^. Everyday function was determined by interview with a significant other, using Reisberg’s IADL-Scale and Global Deterioration Scale, and the Clinical Dementia Rating ^43–45^. Interview evidence from a significant other was not available in 39 PD patients at baseline (only) but we were able to exclude dementia at baseline from a consensus decision derived from contemporaneous clinical test examiner notes and absence of prior dementia during subsequent interview with the significant other. The Montreal Cognitive Assessment (MoCA) was used as a screen for global cognition.

PD-MCI was defined in two ways. First, we applied level II criteria using the most common PD-MCI cut-off for a deficit on any measure, viz. a score at -1.5SD or worse below normative data. Secondly, a logistic regression was fit on all 21 measures simultaneously and, when applied, gave a probability of conversion to PDD. To convert this to a classification, individuals that scored above a threshold on this model were defined as PD-MCI. This threshold could be set to achieve any desired level of sensitivity for detecting conversion to PDD (with a corresponding tradeoff in specificity).To facilitate comparison with the level II criteria, the threshold was set to achieve a similar sensitivity to that determined by the full and reduced batteries (88% and 86%). Models assessing both level II and regression-based PD-MCI definitions used leave-one-out methods to predict the risk of conversion to PDD.

### Statistical analysis

All analyses were conducted in the R statistical environment ^46^. We used a logistic regression model with elastic net regularization, which combines the L_1_ penalty of the Least Absolute Shrinkage and Selection Operator method and L_2_ penalty of the ridge method, to minimise the contribution of non-predictive variables ^47^. This was implemented with the package *glmnet* ^48^. The dependent variable was whether or not conversion to PDD occurred within the 3.5 to 4.5 year period of follow-up. Critically, the evaluation of tests predictive of PDD after baseline testing was conducted independently of any given patient’s scores by using a leave-one-out (LOO) procedure. That is, test selection was performed on N-1 participants, and then the PD-MCI status determined for the Nth individual using the selected tests. This was repeated N times to get an independent PD-MCI status for each individual.

The DRS-2 similarities and ADAS-Cog language measures were introduced after the start of the study period and hence each was missing from 22% of assessments. Picture Completion and Map Search were missing in 4% and 3% respectively. All other tests had <1% missing data. We dealt with missing data using multiple imputation (100 imputed datasets) with a predictive mean matching procedure (the *mice* package) ^49^. To determine the relative risk of PDD arising from having a PD-MCI classification, bootstrapping with 5000 iterations was used for each imputed dataset.

#### Selecting two cognitive measures per domain

Level II PD-MCI criteria require an assessment of at least two tests per five cognitive domains. The impairment cut-off of -1.5SD per measure was based on published normative data or (for action fluency, digit ordering, JLO, and VOSP fragmented letters, similarities component of DRS-2, and language component of ADAS-Cog), local age- and education-adjusted normative data ^17,22,50^. The elastic net model established the two measures most frequently associated with future PDD within each cognitive domain independently (i.e. one model per cognitive domain). To do this, the regularization parameter, lambda, was set in each model such that only two tests predictive of PDD within each domain remained (i.e. the contribution of other tests was forced towards zero within each model). Different pairs of test measures could be selected across the multiple model iterations. The predictors for each of the five cognitive domain models were the z-scores for test measures within that domain. Inclusion frequency across imputation and LOO iterations of a given test was used to provide evidence of whether it was one of the top two predictors in that domain to detect conversion to PDD in the 3.5 to 4.5 year period post-baseline. As it contained only two measures, no analysis was possible for the language domain. The relative risk of PDD for PD-MCI was subsequently estimated on an out-of-sample basis as described in the statistical analysis section.

#### Selecting optimal tests independent of domains

Second, a logistic regression classifier model was used to predict progression to PDD by simultaneously assessing all 21 neuropsychological test measures, disregarding the cognitive domain to which they could be assigned. When all measures in the battery were assessed simultaneously, any measure irrespective of cognitive domain could be associated with progression to PDD. PD-MCI status was defined by the logistic regression classifier score reaching a threshold and required no specific cut-off of test scores (as described in the *PD-MCI and PDD diagnoses* section). Here, the regularization parameter lambda was chosen by 4-fold cross-validation to maximise the AUC plus one standard error ^48^. The model was evaluated out-of-sample (LOO) via AUC. This analysis was also able to examine the influence of cognitive predictors relative to demographic and disease-relevant predictors, namely age, sex, disease duration, years of education, and UPDRS Part-III score.

## Results

There were 145 individuals who remained dementia-free during the duration of the study. There were 51 (26%) participants who converted to PDD. The demographics and neuropsychological test scores of the non-converters and converters to PDD are shown in Table 1. At baseline, the individuals who would subsequently convert to PDD were slightly older, had longer symptom duration, had worse motor symptoms and poorer cognition. Converters to PDD had poorer scores on all neuropsychological test measures, with moderate to large effect sizes.

**Table 1:**
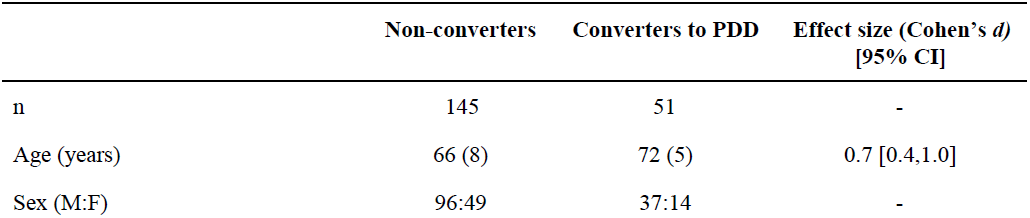
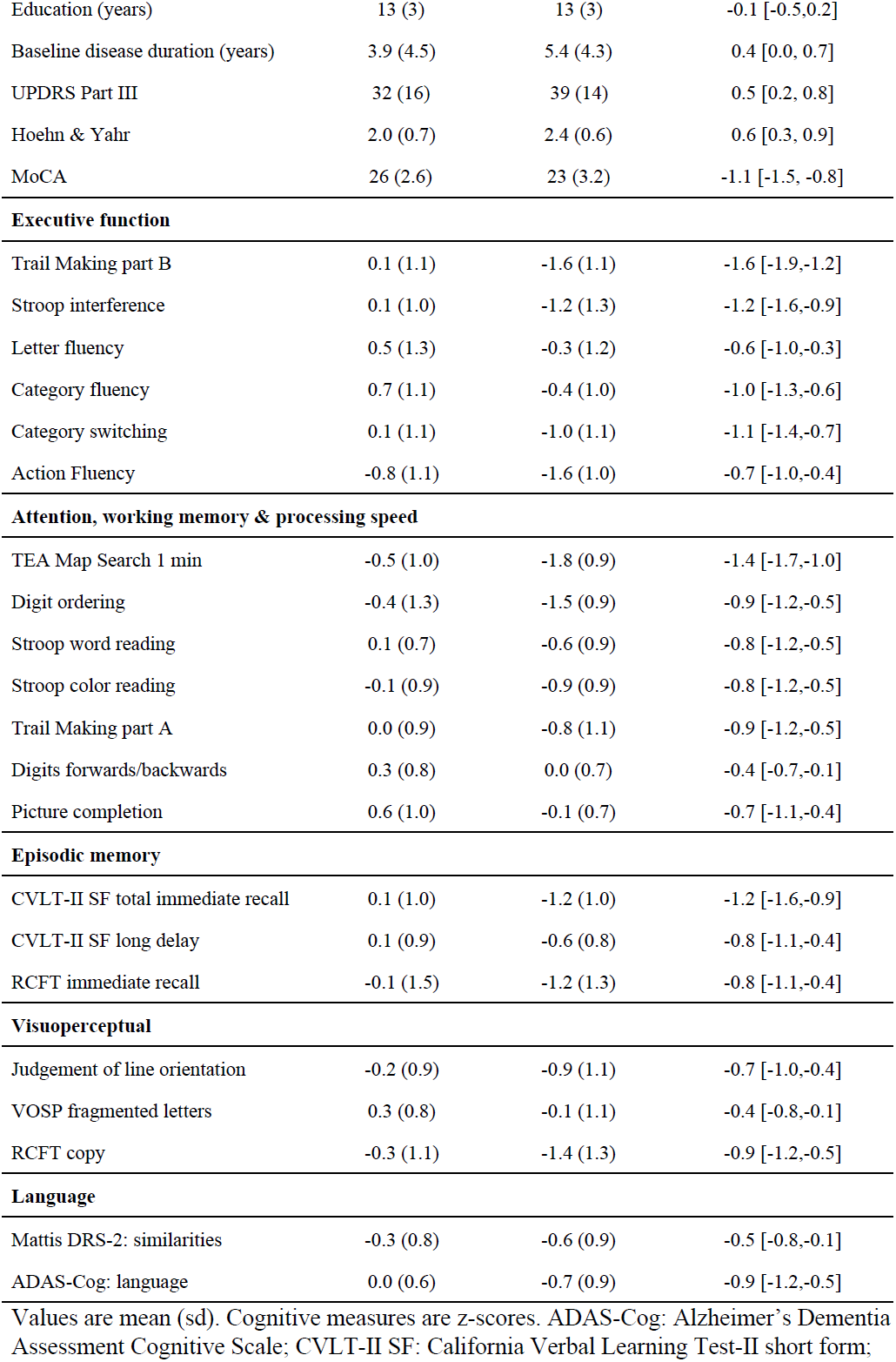
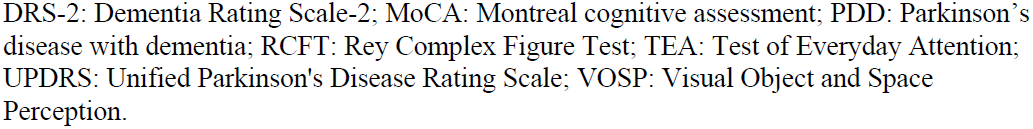
Demographic and cognitive characteristics of the sample at baseline.

|  | Non-converters | Converters to PDD | Effect size (Cohen's *d*)<br>[95% CI] |
| --- | --- | --- | --- |
| n | 145 | 51 | - |
| Age (years) | 66 (8) | 72 (5) | 0.7 [0.4,1.0] |
| Sex (M:F) | 96:49 | 37:14 | - |
| Education (years) | 13 (3) | 13 (3) | -0.1 [-0.5,0.2] |
| Baseline disease duration (years) | 3.9 (4.5) | 5.4 (4.3) | 0.4 [0.0, 0.7] |
| UPDRS Part III | 32 (16) | 39 (14) | 0.5 [0.2, 0.8] |
| Hoehn & Yahr | 2.0 (0.7) | 2.4 (0.6) | 0.6 [0.3, 0.9] |
| MoCA | 26 (2.6) | 23 (3.2) | -1.1 [-1.5, -0.8] |
| **Executive function** |  |  |  |
| Trail Making part B | 0.1 (1.1) | -1.6 (1.1) | -1.6 [-1.9,-1.2] |
| Stroop interference | 0.1 (1.0) | -1.2 (1.3) | -1.2 [-1.6,-0.9] |
| Letter fluency | 0.5 (1.3) | -0.3 (1.2) | -0.6 [-1.0,-0.3] |
| Category fluency | 0.7 (1.1) | -0.4 (1.0) | -1.0 [-1.3,-0.6] |
| Category switching | 0.1 (1.1) | -1.0 (1.1) | -1.1 [-1.4,-0.7] |
| Action Fluency | -0.8 (1.1) | -1.6 (1.0) | -0.7 [-1.0,-0.4] |
| **Attention, working memory & processing speed** |  |  |  |
| TEA Map Search 1 min | -0.5 (1.0) | -1.8 (0.9) | -1.4 [-1.7,-1.0] |
| Digit ordering | -0.4 (1.3) | -1.5 (0.9) | -0.9 [-1.2,-0.5] |
| Stroop word reading | 0.1 (0.7) | -0.6 (0.9) | -0.8 [-1.2,-0.5] |
| Stroop color reading | -0.1 (0.9) | -0.9 (0.9) | -0.8 [-1.2,-0.5] |
| Trail Making part A | 0.0 (0.9) | -0.8 (1.1) | -0.9 [-1.2,-0.5] |
| Digits forwards/backwards | 0.3 (0.8) | 0.0 (0.7) | -0.4 [-0.7,-0.1] |
| Picture completion | 0.6 (1.0) | -0.1 (0.7) | -0.7 [-1.1,-0.4] |
| **Episodic memory** |  |  |  |
| CVLT-II SF total immediate recall | 0.1 (1.0) | -1.2 (1.0) | -1.2 [-1.6,-0.9] |
| CVLT-II SF long delay | 0.1 (0.9) | -0.6 (0.8) | -0.8 [-1.1,-0.4] |
| RCFT immediate recall | -0.1 (1.5) | -1.2 (1.3) | -0.8 [-1.1,-0.4] |
| **Visuoperceptual** |  |  |  |
| Judgement of line orientation | -0.2 (0.9) | -0.9 (1.1) | -0.7 [-1.0,-0.4] |
| VOSP fragmented letters | 0.3 (0.8) | -0.1 (1.1) | -0.4 [-0.8,-0.1] |
| RCFT copy | -0.3 (1.1) | -1.4 (1.3) | -0.9 [-1.2,-0.5] |
| **Language** |  |  |  |
| Mattis DRS-2: similarities | -0.3 (0.8) | -0.6 (0.9) | -0.5 [-0.8,-0.1] |
| ADAS-Cog: language | 0.0 (0.6) | -0.7 (0.9) | -0.9 [-1.2,-0.5] |

Values are mean (sd). Cognitive measures are z-scores. ADAS-Cog: Alzheimer's Dementia Assessment Cognitive Scale; CVLT-II SF: California Verbal Learning Test-II short form;
DRS-2: Dementia Rating Scale-2; MoCA: Montreal cognitive assessment; PDD: Parkinson's disease with dementia; RCFT: Rey Complex Figure Test; TEA: Test of Everyday Attention; UPDRS: Unified Parkinson's Disease Rating Scale; VOSP: Visual Object and Space Perception.

### Selecting two cognitive measures per domain

Figure 2 shows the inclusion frequency for each test within each of the cognitive domains. There was nearly unambiguous selection of two preferred test measures within each cognitive domain over the imputed and LOO datasets. The ten measures across the 5 cognitive domains were: Trail Making part B; Stroop interference; First minute of TEA Map Search; digit ordering; CVLT-II SF total immediate recall; RCFT immediate recall; judgement of line orientation; RCFT copy; Mattis DRS-2 similarities; and ADAS-Cog language.

Table 2 shows the out-of-sample performance metrics of the reduced battery of 10 tests and the performance when all 21 measures were used to identify PD-MCI. The reduced battery classified fewer individuals as PD-MCI, while maintaining a similar sensitivity to capturing converters. As a result, the specificity of the reduced battery increased.

**Table 2:** Performance of the full test battery (21 measures) and two shortened test batteries.

|  | Full 21-test<br>battery<sup>#</sup> | Level-II 10-test<br>battery<sup>*</sup> | Globally optimised<br>battery<sup>*</sup> |
| --- | --- | --- | --- |
| Number classified as PD-MCI /<br>total sample at baseline | 102/196<br>52% | 86.6/196<br>44% | 70.7/196<br>34% |
| PD-MCI conversions to PDD / total<br>conversions to PDD (sensitivity) | 45/51<br>(88%) | 44.0/51<br>(86%) | 44.0/51<br>(86%)<sup>†</sup> |
| PD-N / non-converters to PDD<br>(specificity) | 88/145<br>(61%) | 102.5/145<br>(71%) | 118.3/145<br>(82%) |
| Relative risk of PDD, given<br>PD-MCI [95% CI] | 6.9<br>[3.0, 15] | 8.0<br>[4.3, 24] | 12<br>[6.2, 39] |

<sup>#</sup>Calculated as single estimates from the initial sample; <sup>\*</sup>non-integer values due to the use of the mean over out-of-sample and imputed datasets used to reduce bias from missing data and ensure independence from the test selection procedure; <sup>†</sup>set to this level by choice of threshold for classifier. PD-MCI = PD with mild cognitive impairment. PDD = PD with dementia. PD-N = with cognition in the normal range.

### Selecting optimal tests independent of domains

A mean of 3.3 tests (interquartile range 3-4) were selected as predictors of conversion to PDD across the 19600 model iterations. The tests with the strongest evidence of providing useful predictive information, with inclusion frequencies at or close to 100%, were Trail making part B, TEA Map Search first minute, and CVLT-II SF total immediate recall (Figure 2). Only four other tests were selected in 1% or more of models, which were ADAS-Cog: language (24%), Stroop colour reading (11%), category switching (3%), and RCFT copy (1%). To determine the influence of these tests relative to demographic variables, model fits were extended to include patient age, sex, disease duration, years of education, and UPDRS Part-III score. The overall selection of cognitive tests stayed the same, and only patient age (inclusion frequency 3%) showed any evidence of being weakly predictive in the global model.

Figure 3 (Left Panel) shows the mean coefficient size by any predictor when combined into a global elastic net model. Evidence of a low (or effectively zero) coefficient value in that analysis, however, does not mean the test contained no information relating to PDD conversion. This latter point is illustrated when each test was analysed individually in a GLM. All individual measures had moderate to large coefficient values in terms of an association with conversion to PDD (Figure 3, right panel, all p < 0.05). When analysed separately the demographic variables age, disease duration, diagnosis age, and UPDRS Part-III were also associated with PDD conversion (p <0.05). However, when included in the global elastic net model, they weren’t selected. The out-of-sample global elastic-net logistic regression risk function, applied to each individual, produced an AUC of 0.90 [0.84, 0.96] for identifying converters to PDD vs non-converters. When classifying individuals as PD-MCI or no PD-MCI at baseline, the PD-MCI group had a RR of 12 for conversion to PDD (Table 2).

**Figure 3:**
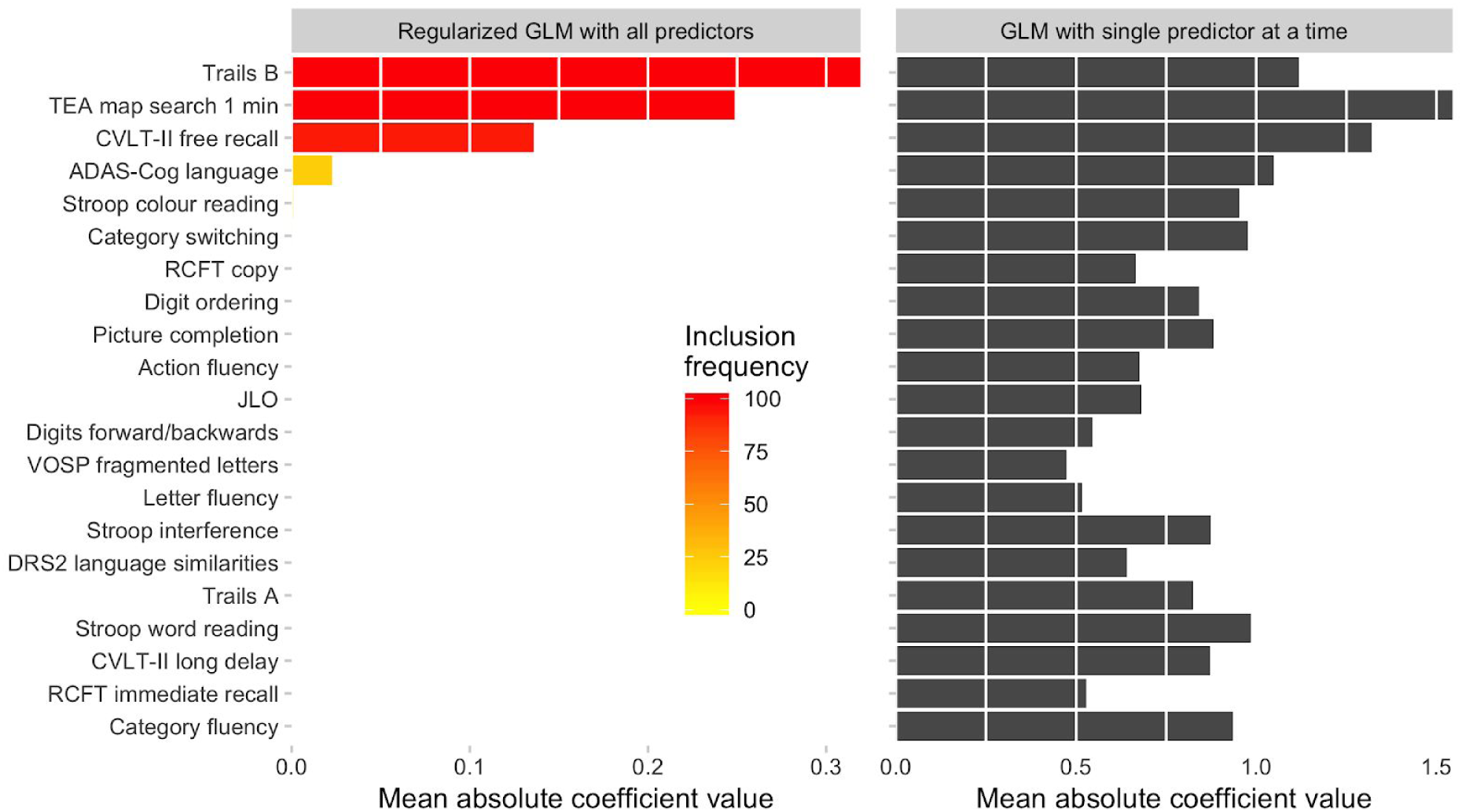
Left: Mean coefficient size per predictor in the global regularized logistic regression function with inclusion frequency (colour); only three tests had strong evidence for inclusion frequency. Right: Coefficient size for each test when analysed in separate GLM models; every test was predictive of progression to PDD when analysed individually.

## Discussion

Three neuropsychological tests showed strong evidence for identifying patients who had an elevated risk of progression to PDD in the next four years. These were Trail Making part B (executive function), Test of Everyday Attention first-minute Map Search (attention), and CVLT-II Short Form total immediate recall (learning and memory). The resulting regression model produced a high AUC (0.90) for detecting conversion to PDD. An impaired cognitive status based upon this regression model resulted in a 12-fold out-of-sample relative risk (lower 95% CI = 6.1). This approach compared favourably with assessments to identify a MDS-TF level II PD-MCI status that used either the whole battery of 21 neuropsychological test measures (RR = 6.9) or 10 test measures, restricted to a pair of tests in each of five cognitive domains (RR = 8.0). Specificity, that is, the percentage who were PD-N (i.e. not meeting PD-MCI criteria) and did not convert to PDD, progressively improved from 61% for the whole test battery to 71% when 10 test measures were selected and 82% when the tests were selected by regression analysis on all 21 measures simultaneously. Across these three analyses, respectively, the positive predictive value for conversion to PDD when a patient was identified as cognitively impaired improved from 44%, to 51% and 62%. Our findings indicate that a 20-minute neuropsychological assessment using Trail Making, Map Search and the CVLT-II Short Form total immediate recall may be sufficient to identify patients at high risk of conversion to PDD in the next four years, and which is not markedly influenced by demographic or motor-related disease characteristics. This would be especially useful when more extensive testing is not feasible.

The reduction from a large battery to two tests per cognitive domain has been proposed when discriminating between level II PD-MCI and PD-N patients ^51^. Here, we show that this approach is also suited with respect to the longitudinal risk of PDD. Beyond the three neuropsychological tests listed above, our analysis added Stroop interference for the executive function domain, digit ordering for attention/working memory, JOL and RCFT-Copy for visuoperceptual function, and RCFT immediate recall for episodic memory. The similarities component of DRS-2 and the language component of the ADAS-Cog were also included for the language domain. One group has already suggested that the initial copy and delayed recall of the RCFT can separate PD-MCI converters from non-converters at baseline ^38,39^. A few studies have reported that poor performance on Trail Making part B, Stroop interference, and JOL are also predictors of progression to PDD, but only Stroop interference has previous evidence of an independent contribution ^27,34–36,39,52,53^. Poor immediate recall of a word list to predict cognitive decline has been reported more frequently ^23,32–36,38,53,54^, but direct comparison with other neuropsychological tests for predicting PDD has been reported only once ^39^. Immediate verbal recall is an episodic memory measure, but poor performance may also reflect impairments in attention or executive control strategies for encoding and retrieval ^29,34^. There has been considerable interest, but mixed evidence, whether poor performance on verbal fluency predicts PDD ^31–36,38,39,55,56^. We found that neither category (semantic) fluency nor letter (phonemic) fluency provided independent evidence for conversion to PDD. No other group has used the Map Search task to predict PDD. Poor performance on Map Search and Trail Making part B may be sensitive predictors because they require a different balance of multiple skills, including complex attention, visuospatial perception, flexibility, motor planning, and speed of processing.

PD-MCI patients who did not convert to PDD within the four-year period are nonetheless probably still at increased risk of conversion in due course ^16,17,24^. The variability in time to dementia conversion highlights the heterogeneity of cognitive impairment and dementia risk across PD patients. The differences between converters and non-converters were not explained by conventional demographic or clinical motor scores. It is likely that phenotypic and genotypic differences, as well as variable neuropathology, comorbidities and lifestyle may contribute to this differential risk ^57–59^. For example, there is strong evidence that other factors we did not measure, such as presence of rapid eye movement sleep behavior disorder or hyposmia, provide independent risk factors for conversion to PDD even in the presence of PD-MCI ^14,60^. Genetic variation, for example in the microtubule-associated protein tau gene, alpha synuclein gene and glucocerebrosidase gene, also influences the risk of dementia ^55,61,62^. Degenerating cholinergic pathways, white matter hyperintensities and cortical thinning are among brain correlates that influence the course of cognitive decline ^63–69^. Another interesting question is whether more fundamental changes in visual perception, including poor visual acuity, contrast sensitivity and color vision, also associate with the risk of dementia ^14,70^.

Potential limitations of our study include the commonly-used cut-off of 1.5SD below normative data to indicate impairment across all tests in our first analysis. There is evidence that -2SD signals patients at greater risk, although this stricter cut-off may increase false negatives ^22,71^. It is also possible that different cut-offs across neuropsychological tests may signal a similar level of risk and this is an inherent outcome in a regression approach. Normative data suffer from problems of comparability across tests due to differences in sensitivity and difficulty. The accuracy for (i.e. confidence in) lower scores in particular may not be equivalent across measures ^72^. The large between-study variability when neuropsychological measures have been used to characterise cognition in PD relative to published norms in the absence of local norms also suggests caution when making comparisons for similar tests across different sites ^21^. A second limitation is that our findings may be test and site-specific. Our sample’s ethnicity was almost entirely of European descent. Replication is needed in other cohorts to confirm the pattern of findings described here and to encourage the establishment of suitable risk scores and eventual translation to the clinic. A third limitation is that the language domain contained only two tests (Mattis DRS-2 similarities component and the composite ADAS-Cog language score). There is mixed evidence whether other language measures such as the full Boston Naming Test or expressive language are appropriate alternatives as predictors of future PDD ^15,16,23,31,35,36,39^. Complex language, such as pragmatic language and comprehension, may be more suitable targets for further study in the context of progression to PDD ^73–75^.

Our study has several strengths that reinforce its value and novelty. Our methods add confidence that the neuropsychological tests selected are sensitive predictors of the risk of PDD over 3.5 to 4.5 years after baseline cognitive assessment. We minimised model overfitting and established out-of-sample predictions for individual patients. The selection was based on a large prospectively-followed sample of established PD patients and we achieved good retention (89%, excluding deceased). The follow-up period used is relevant for capturing significant cognitive decline and conversion to PDD and the proportion of conversion (26%) was similar to that expected from a meta-analysis of prior studies ^15,17–19,24,25,27^. Our study benefited from the use of MDS-TF recommendations for neuropsychological testing and current criteria to confirm PDD status.

We offer two options for patient sample enrichment for therapeutic intervention trials. One abbreviated set of neuropsychological tests satisfies MDS-TF level II requirements (by having two tests per cognitive domain) and affords classification of cognitive subtype. The second, shorter option meets level I requirements (impairment on at least two tests) and potentially yields an increased positive predictive value for conversion to PDD. We substantiated prior findings that Trail Making and immediate verbal recall are valuable indicators of future global cognitive decline in Parkinson’s disease, beyond clinical and demographic variables. We also provide new evidence that Map Search is a particularly sensitive measure of future cognitive decline. This measure of visuospatial attention warrants further study, as we found it to be one of the strongest predictors of conversion from PD-MCI to PDD. A focused set of neuropsychological predictors could be used to improve our understanding of potential biomarkers of disease progression.

## Acknowledgements

We would like to acknowledge individuals who have collected data: Saskia van Stockum, Rachel Nolan, Charlotte Graham, Bob Young, Sophie Grenfell, Krysta Callander, Maddie Pascoe, and Megan Livingstone.

We would like to acknowledge funding from the New Zealand Health Research Council, Brain Research New Zealand—Rangahau Roro Aotearoa, University of Otago, University of Canterbury, Neurological Foundation of New Zealand, Canterbury Medical Research Foundation, and the New Zealand Brain Research Institute. There are no conflicts of interest to disclose.

## Authors’ Roles

1. Research project: A. Conception, B. Organization, C. Execution;
2. Statistical Analysis: A. Design, B. Execution, C. Review and Critique;
3. Manuscript: A. Writing of the first draft, B. Review and Critique.

**DJM**: 1A, 1B, 2A, 2B, 2C, 3A, 3B; **K-LH**: 1C, 3B; **MRM**: 1A, 1B, 1C, 2A, 2C, 3A, 3B; **LL**: 1C, 3B; **TLP**: 1B, 2C, 3B **TRM**: 1B, 2C, 3B; **GJG**: 2C, 3B **TJA**: 1A, 1B, 2C, 3B; **JCD-A**: 1A, 1B, 1C, 2A, 2C, 3A, 3B

## Supplementary Methods

### Details of leave-one-out procedure and bootstrapping

For the LOO procedure, 195 individuals (N-1) were used to establish the selected tests that were predictive of PDD risk using the elastic net regression and then the out-of-sample individual’s status was established as PD-MCI or non-MCI in an unbiased manner based on the tests selected from the in-sample group of 195 patients. The LOO procedure thus initially generated 196 counts of the association of each test measure with progression to PDD and 196 independent tests of the influence of PD-MCI status on progression to PDD.

In this manner, with 100 imputed datasets to take into account the missing data, we had 19,600 iterations (100 × 196) that were used to calculate the inclusion frequency for each test. That is, this frequency was the proportion of iterations in which a given test was selected for inclusion in the generated models.

For each imputed dataset an unbiased relative risk of PDD for PD-MCI vs non-MCI cases could then also be calculated. By resampling each of the 100 datasets (inclusive of imputed data) 5000 times using a bootstrapping procedure, we generated 500,000 estimates to produce a distribution of relative risk for PD-MCI vs non-MCI, which provided the quantification of uncertainty in the relative risk metric.

